# Little pig, little pig, let me come in: The influence of landscape structure and La Niña climatic anomalies on the emergence of Japanese encephalitis virus in Australian piggeries in 2022

**DOI:** 10.1101/2022.12.19.521138

**Authors:** Michael G. Walsh, Cameron Webb, Victoria Brookes

## Abstract

The widespread activity of Japanese encephalitis virus (JEV) in previously unaffected regions of eastern and southern Australia in 2022 represents the most significant local arbovirus emergency in almost 50 years. Japanese encephalitis virus is transmitted by mosquitoes and maintained in wild ardeid birds and amplified in pigs, the latter of which suffer significant reproductive losses as a result of infection. The landscape of JEV outbreak risk in mainland Australia is almost entirely unknown, particularly in the eastern and southern parts of the country where the virus has not been previously documented. Although other areas with endemic JEV circulation in the Indo-Pacific region have demonstrated the importance of wild waterbird-livestock interface in agricultural-wetland mosaics, no such investigation has yet determined the configuration of pathogenic landscapes for Australia. Moreover, the recent emergence in Australia has followed substantial precipitation and temperature anomalies associated with the La Niña phase of the El Niño Southern Oscillation. This study investigated the landscape epidemiology of JEV outbreaks in Australian piggeries recorded between January and April of 2022 to determine the influence of ardeid habitat suitability, hydrogeography, hydrology, land cover and La Niña-associated climate anomalies in demarcating risk. Outbreaks of JEV in domestic pigs were associated with ardeid species richness, agricultural and riparian landscape mosaics, hydrological flow accumulation, and grasslands. This study has identified the composition and configuration of landscape features that delineated risk for piggeries during the 2022 emergence of JEV in Australia. Although preliminary, these findings can inform actionable strategies for the development of new One Health JEV surveillance specific to the needs of Australia.

## Introduction

Japanese encephalitis virus (JEV) re-emerged in Australia over the 2021-2022 summer with extensive outbreaks in piggeries and sporadic cases in humans manifesting an unprecedented geographic distribution across Queensland, New South Wales (NSW), Victoria, and South Australia[1–3]. Previously, documented JEV circulation had been limited to very localised areas in the Torres Strait Islands[4] and the Cape York Peninsula of northern Queensland[5], but with evidence of regional ongoing circulation between southern Papua New Guinea, the Torres Strait Islands, and northern Australia[6]. The extent of these outbreaks indicates an expanded circulation of JEV much further south than has previously been recognised or predicted[6]. This re-emergence and redistribution of JEV in Australia is of considerable concern given the impact on the domestic pig industry and the severe human health consequences among those who present with clinical disease. Infection with JEV can cause severe reproductive losses in pigs, as well as concern for spillover to humans[7]. In clinically detected human cases, estimated case fatality rates range from 14—30% and of those who survive, an estimated 49% suffer permanent neurological sequelae including ‘locked-in syndrome’[8]. However, the epidemiology of human infection is difficult to determine because most infections are asymptomatic, with less than 1% of infections presenting with clinical disease[9]. As such, cryptic spillover of JEV to humans from competent maintenance or amplification hosts is the norm rather than the exception.

Japanese encephalitis virus is a mosquito-borne zoonotic virus that circulates in wild ardeid bird maintenance hosts[10–13], and is frequently amplified in domestic pig hosts[14–19]. In Australia, evidence suggests that *Culex annulirostris* is the primary vector of JEV[20], although there is some evidence to suggest additional invasive species may also be relevant in some landscapes, such as *Cx. tritaeniorhynchus* and *Cx. gelidus*, both of which are well-established JEV vectors across the Indo-Pacific region, and *Cx. quinquefasciatus*, which is well established throughout Australia[21]. There remains much to learn about the role of local mosquito species, their abundance, and diversity in driving transmission and spillover in Australia. However, it is likely that the ecological niches of key species must be carefully considered[22]. Previous work investigating the landscape epidemiology of JEV across endemic areas of the Indo-Pacific region, have shown that anthropogenic ecotones between cultivated land and riparian and other freshwater marsh wetlands are associated with considerable risk of outbreaks[23]. In contrast, we have very little understanding of the nature and distribution of risk across landscapes within Australia given the very low occurrence and limited geographic spread of JEV outbreaks prior to 2022. Given the recognised importance of maintenance and amplification hosts to both endemic and epidemic JEV transmission, similar environmental features that favour these host-pathogen transmission cycles throughout Asia may be anticipated to influence risk in Australia. However, landscape composition and configuration in relation to JEV outbreak risk has yet to be investigated in Australia, where alternative abiotic or biotic elements may feature more prominently in determining risk. For example, the widespread distribution of feral pigs in Australia, and their potential interaction with waterbirds and mosquitoes, may present pathogenic landscapes unique to the continent.

The recent emergence of JEV in Australia coincided with a period of two La Niña phases of the El Niño Southern Oscillation (ENSO). The La Niña phase of ENSO is characterised by a shift in ocean currents that leads to a build-up of warmer than average surface waters in the western Pacific Ocean, which leads to greater than average precipitation and lower than average temperatures in eastern Australia[24]. Accordingly, La Niña contributed to extensive precipitation and temperature anomalies throughout 2021 across much of eastern Australia[25]. Given the sensitivity of vector mosquitoes and reservoir maintenance hosts to climate, anomalous La Niña precipitation and temperature patterns may also have contributed to the emergence and wide distribution of JEV across eastern Australia in 2022, although there is currently no conclusive evidence regarding the pathway or timing of this recent JEV introduction[6]. Similar trends in spread and activity of vector-borne flaviviruses such as West Nile (Kunjin subtype) virus and Murray Valley encephalitis virus are generally associated with above average rainfall occurring in conjunction with La Niña influence[26,27]. Indeed, climate anomalies may be particularly relevant for temperate eastern Australia, which does not typically experience substantial seasonal extremes in precipitation. The typical climate pattern of temperate eastern Australia is in stark contrast with other JEV endemic areas in the region, such as India, where the marked seasonal extremes of precipitation and temperature associated with the South Asian monsoon is a critical driver of seasonal JEV outbreaks[28].

The objective of the current investigation was to identify key indicators of JEV outbreak risk to piggeries across eastern Australia by interrogating surrounding landscape characteristics. Specifically, this study explored the extent to which wildlife-host habitat suitability, proximity to wetlands and other land cover, hydrological geomorphology, and climate anomalies associated with the 2021 La Niña were associated with piggery outbreaks of JEV. It was anticipated that outbreaks in piggeries would be associated with greater ardeid suitability, increased proximity to wetlands and cultivated land, increased La Niña-associated precipitation, and increased hydrological flow accumulation.

## Materials and Methods

### Data sources

#### Animal data

Fifty-four location-unique piggery outbreaks were reported to the World Organisation for Animal Health (WOAH) by the Australian government between 19 January, 2022 and 4 April, 2022[29].Outbreak is defined here as the occurrence of one or more cases at the level of the piggery. These 54 outbreaks were used to train the models (described below) after verifying the geographic coordinates of each location in Google Maps. In addition to the 54 outbreaks, we acquired an additional 11 independently-documented locations positive for JEV (8 additional piggeries reported by a commercial pork producer, and 3 positive mosquito pools that were detected by the NSW Arbovirus Surveillance and Mosquito Monitoring Program[30]). These 11 locations were used as an external validation of model performance as described in the analysis section below.

There are 14 extant species of ardeid birds in mainland Australia[31]. A total of 791,416 observations of these species recorded between 1 January, 2010 and 31 December, 2020 were obtained from the Global Biodiversity Information Facility (GBIF)[32] to model the landscape suitability of each species and to generate a proxy for species richness across eastern Australia. Similarly, 9,667 observations of feral pigs recorded over the same time period were also obtained from the GBIF[33] to model feral pig suitability. Domestic pig density and piggery density data were acquired from the national herd dataset that is used in the Australian Animal Disease Spread Model, in which > 8000 registered pig herds of all types (including commercial, boar producers, smallholder and pig keepers) are recorded[34].

Reporting bias may influence the recording of wild bird observations, particularly with respect to differential reporting in regional and remote locations. Therefore, this study corrected for such bias by selecting background points proportional to the human footprint (HFP), which is a robust indicator of landscape accessibility. The HFP data product has been described in detail[35]. Briefly, HFP was constructed based on population density, rural versus urban location, land cover, artificial light at night, and proximity to roads, rail lines, navigable rivers, and coastline. These items were scored and summed to calculate the human influence index (HII). This index ranges from 0 (signifying no human impact) to 64 (signifying maximum human impact). The HFP is then calculated as the ratio of the range of HII values in the local terrestrial biome to the range of HII values across all biomes and is expressed as a percentage[35]. The HFP data were acquired as a raster from the Socioeconomic Data and Applications Center (SEDAC) registry maintained by the Center for International Earth Science Information Network (CIESIN) [36]. The reporting of piggery outbreaks may also be affected by bias, with commercial herds, especially larger herds, being more likely to observe and report cases due to systematic recording of production data. As such, we further corrected for piggery reporting bias by selecting background points for JEV models proportional to herd size (see the statistical analysis section below).

#### Environmental data

All wetlands[37,38] and land cover data[39] were obtained as 3 arc second data products from the European Space Agency and Climate Change Initiative, which are archived with the WorldPop data hub[40]. Two distinct time points of remotely sensed land cover were obtained for the two analyses described below. Land cover assessed in 2010 was applied to the ardeid species distribution models, as this time point corresponded to the beginning of the period of recorded bird observations described above, whereas the latest available land cover assessment (2015) was applied to the JEV models to reflect the time point most proximal to the outbreaks (see the statistical analysis description below). A separate high-resolution (3 arc seconds) raster data product for all rivers and waterways produced in collaboration between CIESIN and the WorldPop project was also acquired[41]. Hydrological flow accumulation quantifies the amount of upland area draining into each 500m X 500m area, and thus is a metric for water movement through, and accumulation in, the landscape. This data product was obtained from the Hydrological Data and Maps based on SHuttle Elevation Derivatives at multiple Scales (HydroSHEDS) information system[42].

Weather anomaly data were obtained from the Goddard Earth Science Data Information and Services Center[43] and comprised the 2021 mean monthly precipitation, temperature, and soil moisture anomalies. Each of the 3 measurements represents the difference between each month of 2021 and the baseline measurement recorded for that month over the period 1982 to 2016. Precipitation was recorded as rainfall flux (kg/m^2^/s) and was converted to millimetres per day for analysis. This dataset thus provides a monthly record of change in precipitation, temperature, and soil moisture from the climate baseline under the La Niña phase conditions experienced throughout the year prior to the emergence of JEV in piggeries in early 2022. WorldClim was used as the source of baseline climate data[44] for modelling ardeid habitat suitability and comprised the mean annual precipitation, mean annual temperature, and isothermality.

The Priestley-Taylor α coefficient (P-Tα) is the ratio of actual evapotranspiration to potential evapotranspiration and was used to quantify water stress in the landscape for the modelling of ardeid habitat suitability[45,46]. The P-Tα represents the water availability in the soil and the water requirements of the local vegetation, contextualised by solar energy input. The P-Tα raster data were obtained from the Consultative Group for International Agricultural Research (CGIAR) Consortium for Spatial Information at a resolution of 30 arc seconds[47].

### Statistical analysis

#### Ardeidae habitat suitability modelling

The habitat suitability of each of the 14 ardeid species extant in Australia, and feral pigs, was modelled using an ensemble of three species distribution modelling (SDM) frameworks: random forest (RF), boosted regression trees (BRT), and generalised additive models (GAM). The two former approaches (BRT and RF) employ machine learning frameworks that algorithmically optimise homogeneity among a response (e.g. species presence) and a set of environmental features. The optimised decision trees can capture complex interactions between predictors[48–51]. In contrast, GAMs fit multiple basis functions with smoothed covariates thus allowing for the fitting of nonlinear relationships between species presence and environmental features[52,53]. Each habitat suitability model under the three distinct modelling frameworks (BRT, RF, and GAM) applied 5-fold cross-validation. Species presence data were thinned to include only one observation per pixel in the analysis to prevent overfitting (Supplementary Table S1). Distance to all inland wetlands, forest, shrubland, herbaceous vegetation (this is predominantly grassland land cover in subtropical and temperate eastern Australia, and is hereafter referred to as grasslands), aquatic vegetation, and cultivated land cover, P-T alpha, isothermality, mean annual temperature, and mean annual precipitation, comprised the environmental features included in the habitat suitability models. The environmental features exhibited low correlation overall (all Pearson’s correlation coefficients < 0.5). In addition, the variance inflation factor was < 8 for each feature included in any suitability model. Therefore, collinearity was not a concern for the fitted models. Each of the three habitat suitability model frameworks (BRT, RF, and GAM) was evaluated for fit and performance for each ardeid species. Model fit was assessed via the deviance, while model performance was assessed via the area under the receiver operating characteristic curve (AUC). An ensemble estimate of habitat suitability was then produced for each species from the three suitability frameworks using their weighted mean (weighting based on AUC)[54]. Background points used in the habitat suitability models were sampled proportional to the human footprint to correct for potential spatial sampling bias among the GBIF occurrences. Species habitat suitability was modelled at a scale of 30 arc seconds (∼1 km). Each species is presented in Supplementary Table S1 with their corresponding number of field observations, thinned analytical observations, and model metrics.

After modelling the distributions of individual Ardeidae species’ landscape suitability, a stacked composite of ardeid suitability was summed across all individual species suitability distributions as a proxy for regional species richness. It is important to note that the construct of species richness used for the current study is not intended to represent local community composition, and therefore is not a true measure of local species richness. Local community structure cannot be adequately measured without accounting for interspecific interaction (particularly competition in the ardeid context) and dispersal ability, both of which would be expected to influence local community assembly[55]. However, this metric does provide utility at regional scale, where environmental filtering would be expected to be a key driver in determining the composition of the regional species pool[55]. As the estimates of individual species habitat suitability represent the potential for environmental filtering based on species’ niches[55], we thus consider the stacked suitabilities as a useful proxy for the regional species pool rather than local species richness. As such, the specific interpretation of ardeid richness in the context of the scale of analysis under the current investigation, is the number of species from the regional pool that are capable of colonising suitable patches and contributing to local community assembly. Realised local community assembly, however, will also be determined by interspecific interaction and dispersal ability. The sdm package[54] for the R statistical software platform, version 4.1.2 [56], was used to fit each model and to derive the three-model ensembles for each species.

#### Point process modelling

Outbreaks of JEV in piggeries were modelled as a point process using inhomogeneous Poisson models[57]. These models allow the evaluation of spatial dependencies among JEV outbreaks in relation to landscape features and thus help to spatially demarcate risk. As a null model representing complete spatial randomness (CSR), JEV outbreaks were first fitted as a homogeneous Poisson process, with conditional intensity,

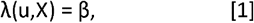

in which u represents outbreak (X) locations and β is the intensity parameter which signifies the number of points in a subregion of a defined spatial window. The expected intensity under CSR is simply proportional to the area of the subregion under investigation[57], indicating an absence of spatial dependency.

The null CSR model was compared to an inhomogeneous Poisson process, which incorporates spatial dependency for JEV outbreak occurrences into the model structure and has conditional intensity,

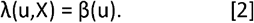

Here outbreak intensity is modelled as a function of outbreak location, u. Since spatial dependence in JEV outbreaks was indicated (see results below), simple and multiple inhomogeneous Poisson models were fitted with landscape feature covariates as follows:

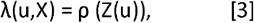

in which ρ represents the association between the JEV outbreak intensity and landscape feature Z at location u. Associations between outbreaks and landscape features were computed as relative risks from the model regression coefficients.

Environmental covariates were aggregated to 5 arc minutes, the minimum resolution to which all reported outbreaks could be reliably located. Since outbreaks were enumerated at the level of piggeries, piggery density was used as the offset in all the point process models so that these models appropriately represented risk to pigs. As described above, background points were sampled proportional to mean herd density (per 5 arc minutes of resolution) to correct for potential outbreak reporting bias. We note that piggery density and mean herd density were not correlated (Pearson’s correlation coefficient = 0.06) and so background points were not sampled proportional to the same or similar feature that serves as the offset in the point process models. The crude associations between JEV outbreaks and distances to inland wetland and riparian wetland systems, distances to forest, shrubland, grassland, aquatic vegetation, and cultivated land cover, hydrological flow accumulation, regional Ardeidae richness, feral pig suitability, and La Niña precipitation, temperature and soil moisture anomalies were each assessed individually with a separate simple inhomogeneous Poisson model (Supplementary Table S2). A composite of temperature anomaly was computed as the mean annual temperature below the climate average in degrees Celsius across all months in 2021. Two composite measures of precipitation anomalies were computed. First, the mean annual precipitation above the climate average in millimetres per day was computed across all months. Second, the mean precipitation above the climate average in millimetres per day averaged between June and September, which are two months that receive some of the lowest precipitation across much of temperate and subtropical eastern Australia over the climate average and were also the only individual months that demonstrated an association between positive precipitation anomalies and JEV risk (Supplementary Figure S1, Supplementary Table S3). Those landscape features univariably associated with outbreaks were included as covariates in the multiple inhomogeneous Poisson models (Supplementary Table S2, Supplementary Figure S2, Supplementary Figure S3). The features included as covariates in the multiple inhomogeneous Poisson models demonstrated low correlation (all values of the Pearson’s correlation coefficient were ≤ 0.53) and low variance inflation factors (all VIF ≤ 4.07) and were therefore considered appropriate for inclusion together in the point process models. Effect modification between riparian systems and cultivated land was additionally considered by including an interaction term in the best-fitting model. The point process models were assessed according to fit, using the Akaike information criterion (AIC), and performance, using the AUC. Furthermore, an independent dataset based on additional piggery outbreak reporting and mosquito and sentinel chicken surveillance (as described above) was used to test model performance, thus providing a crude test of external validity. The full model was compared to reduced models nested on four domains of landscape features (hydrogeography, land cover, ardeid hosts, and climate) to determine model selection. These models were also compared to a model derived from a stepwise selection procedure with the full point process model[58,59]. To identify whether the landscape features in the final model accounted for the spatial dependencies observed in JEV outbreaks, K-functions fitted to the outbreaks before and after point process modelling were compared. The R statistical software version 4.1.2 was used to perform the analyses[56]. Point process models were fitted, and K-functions estimated, using the spatstat package[58,59].

## Results

The distribution of the 54 piggery outbreaks across eastern Australia along with their kernel density estimate is presented in Figure 1. The model with the best fit and performance (Model 1, Table 1; Supplementary Table S4) demonstrated strong associations between JEV outbreaks and ardeid richness (RR = 3.43; 95% C.I. 1.317 – 8.917), proximity to waterways (RR = 0.93; 95% C.I. 0.862 – 0.997), cultivated landscapes (RR = 0.73; 95% C.I. 0.650 – 0.823), and hydrological flow accumulation (RR = 1.002; 95% C.I. 1.0004 – 1.003). Increased La Niña-associated precipitation and decreased temperature were both associated with increased risk univariably, but these associations did not persist after accounting for landscape structure and ardeid suitability. Of note, there was considerable increased precipitation across eastern Australia, but these anomalies in precipitation manifested extensive geographic heterogeneity from month to month and were associated with JEV outbreaks only for those of months of the year that historically receive the lowest rainfall (Supplementary Figure S1; Supplementary Table S3). Importantly, the functional form of the association between JEV outbreaks and ardeid richness was quadratic (Table 1), such that outbreak risk increased as richness increased from 0 to 6 ardeid species, but then dropped off as richness increased still further from 7 to 14 species. In addition, note that proximity to waterways and cultivated land is quantified by distance to these features such that the inverse relative risks indicate decreasing outbreak risk with increasing distance from each feature and vice versa. The distribution of piggery risk based on this model is presented in Figure 2 along with the 95% confidence limits for the estimate. The region of greatest risk extends westward from the Great Dividing Range to the south coast of Victoria and the coast of south Australia. To more thoroughly interrogate the uncertainty associated with risk projection particularly given the small sample size, a sensitivity analysis was conducted to evaluate model performance at the upper and lower limits of the outbreak risk estimate. Model performance at the upper confidence limit (AUC = 93.8%) was very similar to model performance based on the risk estimate (AUC = 93.5%), while performance at the lower confidence limit was moderately diminished (AUC = 88.0%) but nevertheless still good. The configuration of landscape features adequately explained the spatial dependency manifested among the JEV outbreaks as demonstrated by comparing the homogeneous and inhomogeneous K-functions (Figure 3).

**Table 1.**
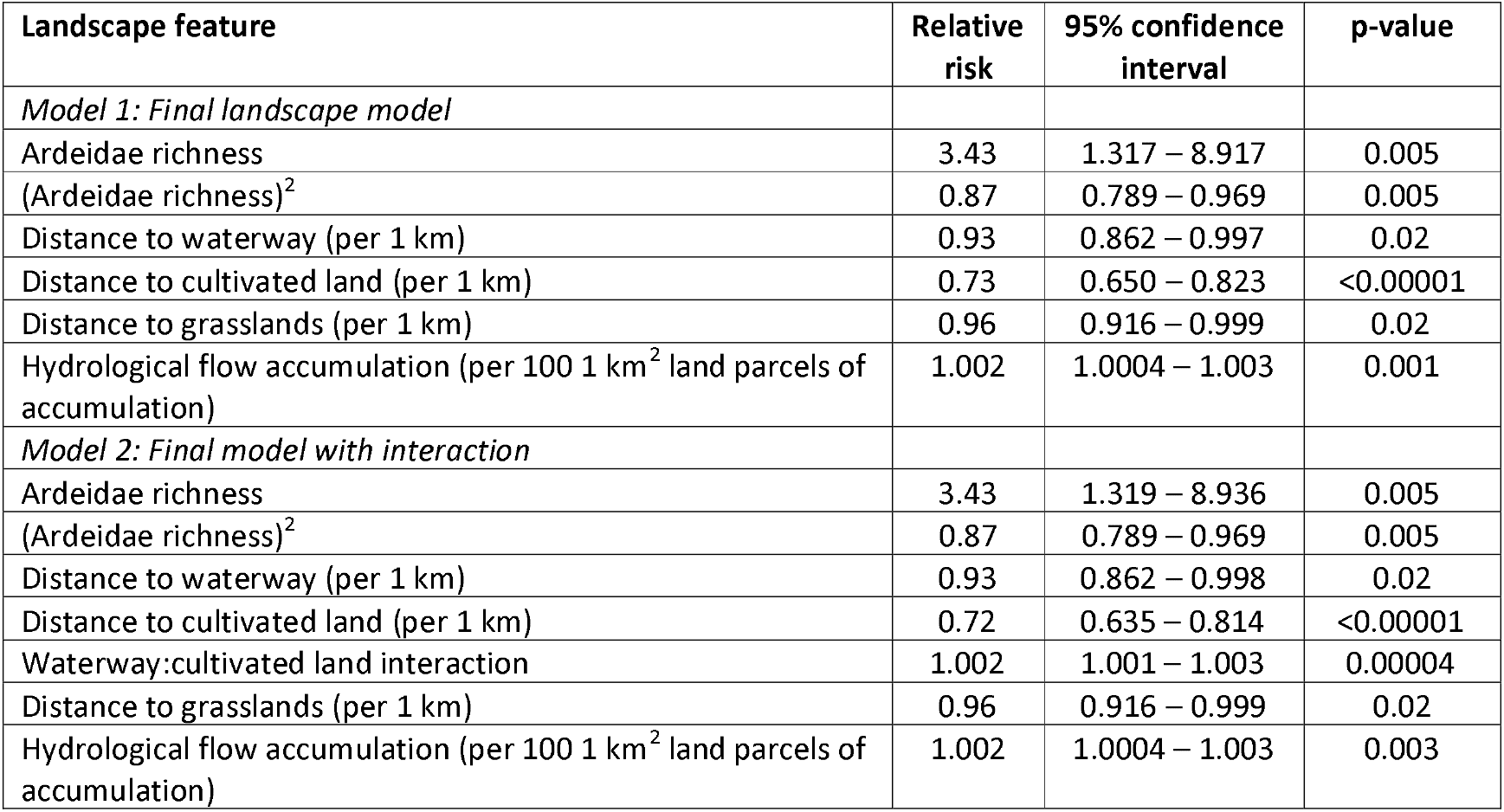
Adjusted relative risks and 95% confidence intervals for the associations between Japanese encephalitis virus (JEV) outbreaks and each landscape feature as derived from the best fitting inhomogeneous Poisson model. Each landscape feature is adjusted for all others in each of the two models.

**Figure 1.**
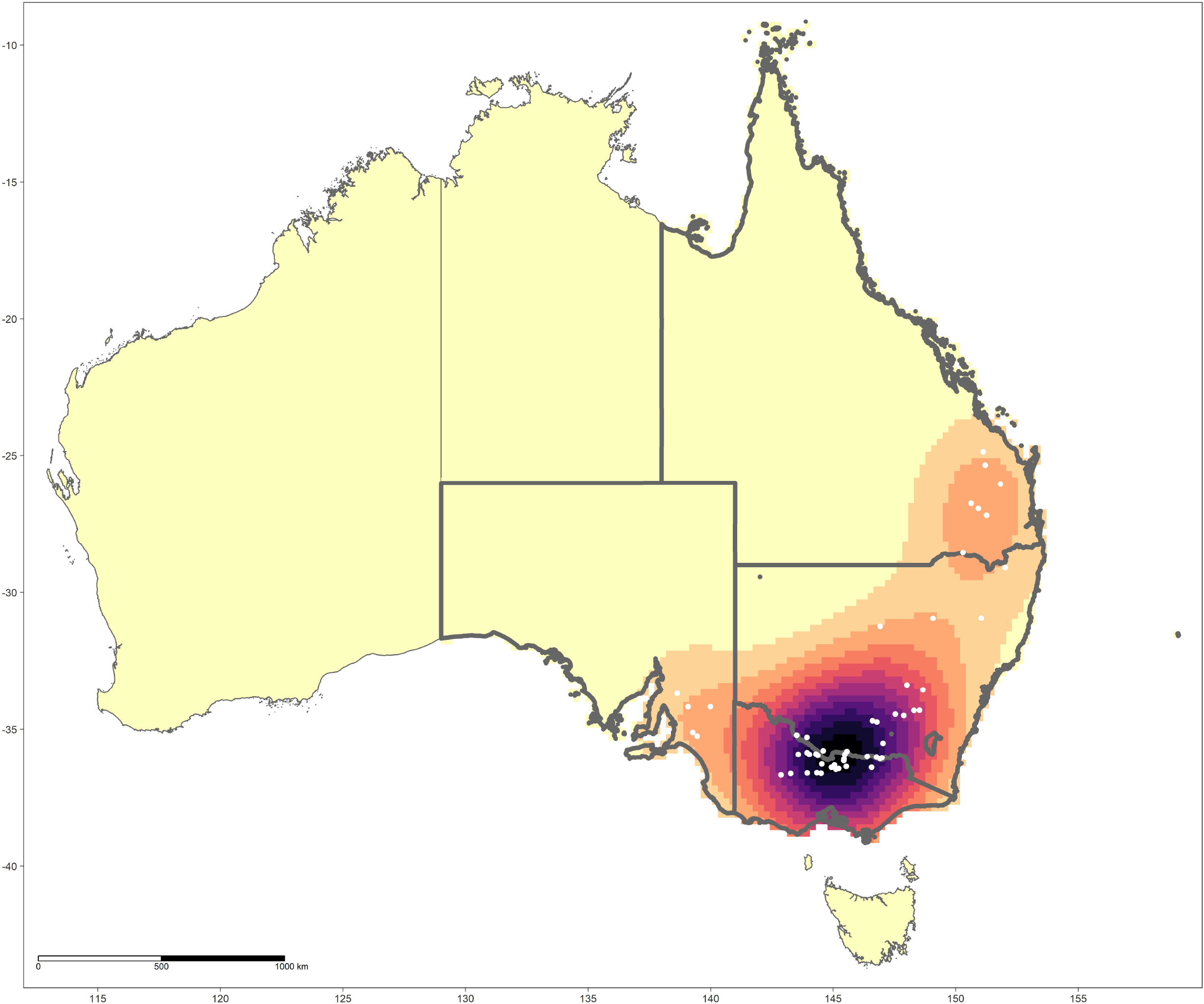
The distribution of Japanese encephalitis virus (JEV) outbreaks in piggeries across eastern and southern Australia and their kernel density estimate. Affected states are highlighted with borders in bold.

**Figure 2.**
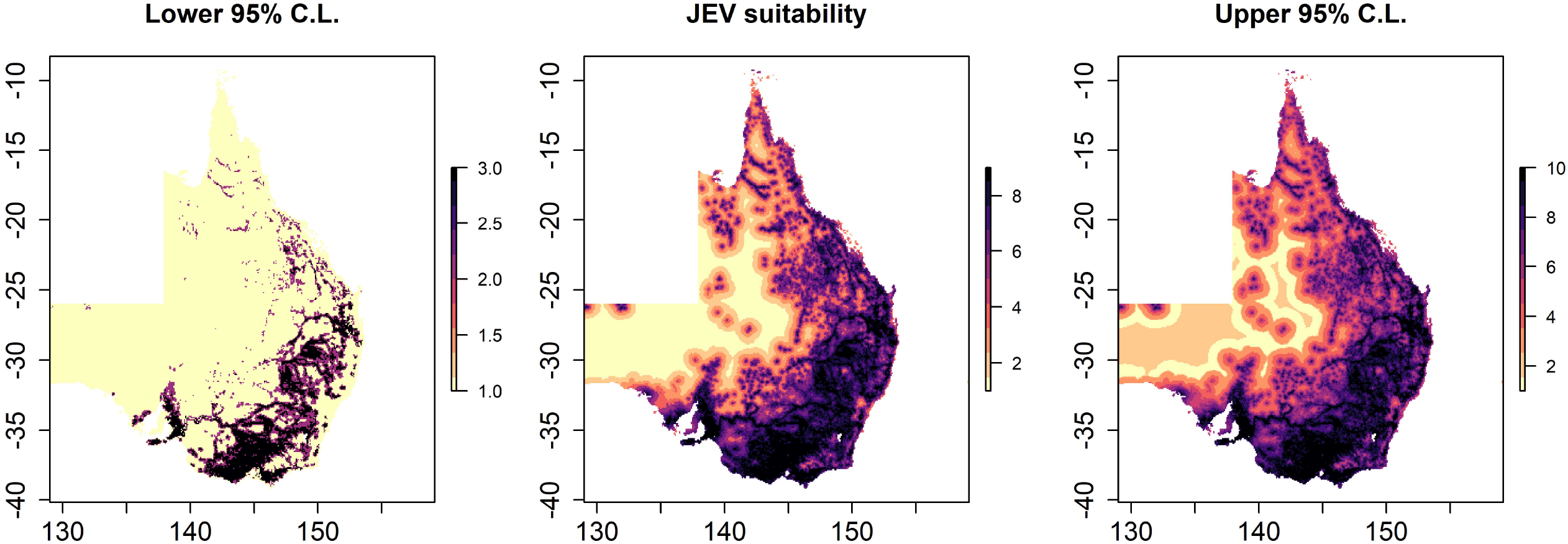
Japanese encephalitis virus (JEV) outbreak risk based on intensity estimates at 5.0 arc minutes. The distribution of JEV predicted intensities is presented in the centre panel, while the left and right panels present the lower and upper 95% confidence limits for the predicted intensities, respectively. Predictions are based on the best fitting and performing inhomogeneous Poisson point process model (Table 1, Model 1).

**Figure 3.**
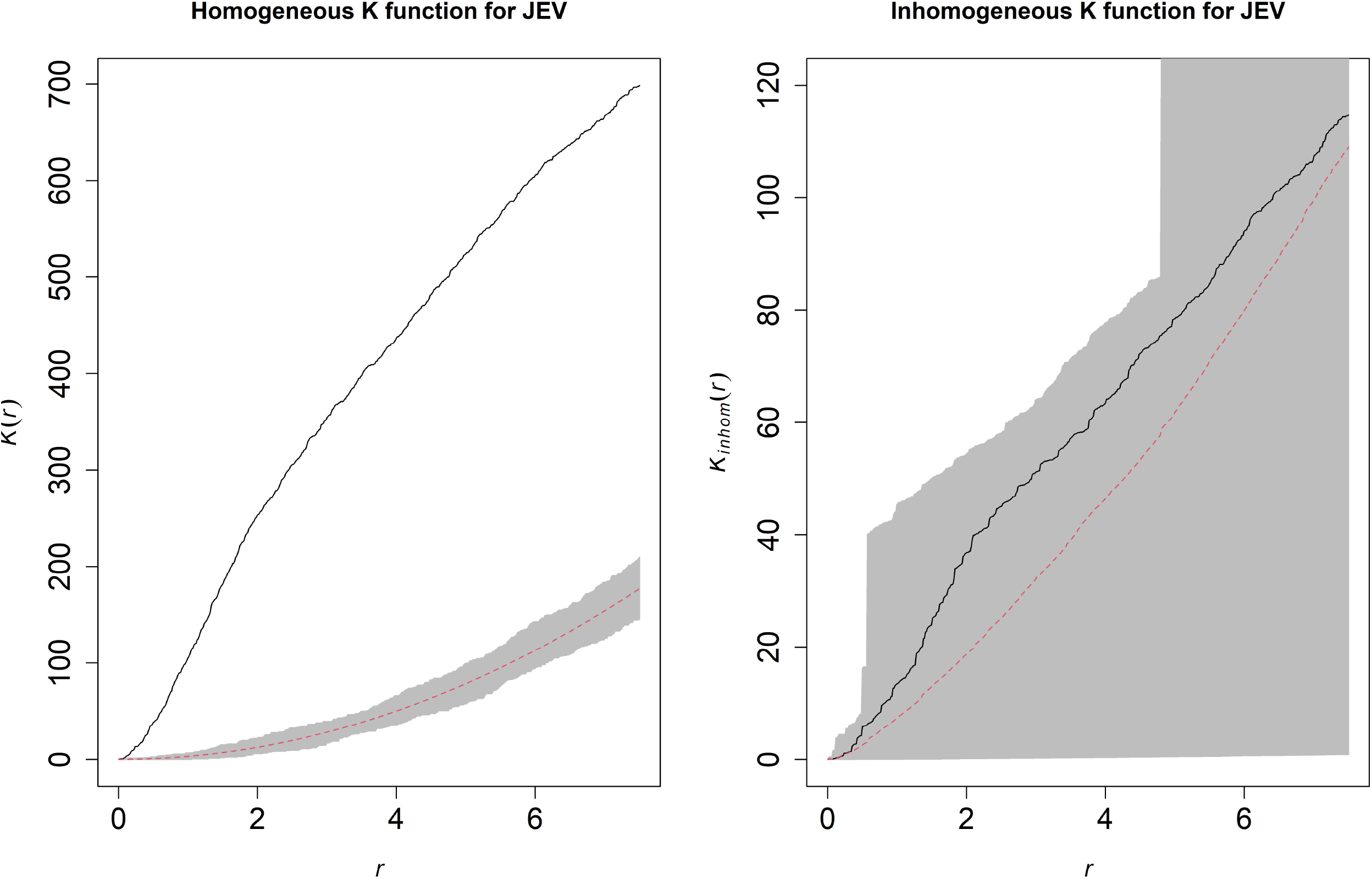
Estimated homogeneous (left panel) and inhomogeneous (right panel) K-functions for Japanese encephalitis virus (JEV) outbreaks. The homogeneous K-function is not an appropriate fit due to the spatial dependency in JEV outbreaks as depicted by the divergent empirical (solid line) and theoretical functions (the latter is the theoretical function under complete spatial randomness, represented by the dashed line with confidence bands in grey). Conversely, the inhomogeneous K-function indicates that the model covariates sufficiently accounted for the spatial dependency (overlapping empirical and theoretical functions). The x-axes, r, represent increasing radii of subregions of the window of JEV outbreaks, while the y-axes represent the K-functions.

Because several JEV endemic settings across the Indo-Pacific region have shown a strong association between JEV risk and the sharing of space between wetland habitat and cultivated land, this relationship was further explored in the current study despite the model with effect modification demonstrating a slightly poorer fit (Model 2, Table 1; Supplementary Table S4). As anticipated, this model demonstrated that proximity to both waterways (RR = 0.93; 95% C.I. 0.862 – 0.998) and cultivated landscapes (RR = 0.72; 95% C.I. 0.635 – 0.814) was associated with increased risk. However, risk was most pronounced at locations proximal to both (interaction term RR = 1.002; 95% C.I. 1.001 – 1.003), whereby increasing distance from one, even while remaining proximal to the other, was associated with diminishing risk.

## Discussion

This is the first report describing the landscape epidemiology of JEV in Australia and the specific threat posed to piggeries. It represents an important initial step for the development of a One Health surveillance system to protect the health of both Australian livestock and humans. Risk was most pronounced in landscape mosaics of riparian systems and agriculture, which demonstrated a high degree of water flow accumulation. In addition, intermediate regional ardeid richness was associated with JEV risk, wherein habitat suitable for a community assemblage of intermediate richness was marked by high risk while habitats suitable for low or high potential community assemblages were marked by lower risk. Finally, the associations between JEV risk and La Niña climate anomalies, including increased precipitation and decreased temperature, were weaker than anticipated.

Herons and bitterns (Ardeidae family) are key maintenance hosts for JEV [10–13], while pigs are recognised as important amplifying hosts [14–19]. As such, transmission between waterbirds and mosquitoes poses a risk of spillover to local piggeries that may subsequently increase the risk of ongoing transmission. The current study reinforced previously identified associations with ardeid habitat in the Indo-Pacific region[23] and suggests that waterbird arbovirus sampling should be implemented as a priority for JEV surveillance, particularly in landscape mosaics of riparian wetlands and agriculture. It is acknowledged that surveillance of JEV in waterbird populations is operationally challenging, but valuable insights for animal and human health could be gained if the challenges can be met. An important remaining question concerns the qualitative extent to which waterbirds share patches with domesticated pigs and mosquito vectors in landscapes comprised of both wetland habitat and agricultural land use. Currently we do not know if JEV transmission dynamics are strongly influenced by interspecific interaction among the vertebrate hosts, for example by way of local community composition and the dilution effect[60], or if conduits to transmission are primarily opened by the behaviour of the vectors, or some combination of both. Moreover, it is unclear if there are maintenance host species that are particularly important for the dispersal and local transmission of JEV in Australian landscapes, and to what extent such species may or may not be synanthropic generalists. While exploration of the contribution of individual species to local community JEV transmission is beyond the scope of the current study due to the scale of outbreak reporting, it is worth noting that four out of the six previously documented competent ardeid host species, *Egretta garzetta*[11,61,62], *Egretta intermedia*[12,61,62], *Ardea alba*[63], *Bubulcus coromandus*[10,63] are all synanthropic and widely distributed throughout eastern Australia.

The association between riparian wetlands and JEV outbreaks is intuitive since these systems provide important habitat for both the ardeid waterbird hosts described above and key vector mosquitoes, particularly *Cx. annulirostris*, which favours diverse freshwater wetland habitats[64], and is believed to be the primary vector of JEV in Australia[20]. Interestingly, this specific association with riparian wetlands has been previously identified with JEV risk in other areas throughout the Indo-Pacific region[23]. Importantly, the greatest risk was not associated with pristine riparian systems, but rather with riparian systems that share landscapes with agricultural land, thus highlighting the importance of habitat fragmentation and emergent anthropogenic ecotones to risk. There may be a suppressive influence on mosquito abundance in areas of higher diversity of potential mosquito predators, which are likely to be greater in less disturbed landscapes[65,66]. As expected, landscape patches receiving more upland surface water accumulation were at higher risk of JEV outbreaks, which fits with the findings of riparian proximity since these systems generally receive the highest surges of runoff following rainfall and indicates that landscapes prone to flooding may require enhanced monitoring for JEV to improve current surveillance systems.

The relatively weak associations with La Niña-associated weather anomalies, particularly increased precipitation, was somewhat surprising. Interestingly, areas receiving higher than average rainfall in the months that typically receive the lowest rainfall throughout the year (June-September) in temperate eastern Australia were univariably associated with increased JEV outbreaks. However, this association did not persist after accounting for landscape structure and ardeid suitability. Interpretation of this finding requires caution. The extent to which precipitation and temperature anomalies influence viral circulation among vectors and hosts is likely to be scale-dependent. For example, temperature and precipitation influence vector population dynamics and mosquito biology (e.g. host-seeking and longevity) as well as viral ecology (e.g. extrinsic incubation rates) on a local scale, which could not be captured under the current analysis.

All La Niña-associated weather anomalies were assessed cumulatively in this study, so the evaluation of direct effects of specific time-lagged weather events was not possible. Nevertheless, it was anticipated that the extensive precipitation anomalies, in particular, may leave a substantial footprint across the landscape with respect to risk. However, given the relative broad-scale of the current study, the impact of increased La Niña precipitation may be dilute, but not sufficiently dilute to preclude the univariable association. However, after incorporating additional landscape features that manifest a footprint that is broader in scale, the association with increased precipitation did not persist. It is also possible that the effects of precipitation may be direct, but manifest in locations that are far removed from where the heaviest rainfall was experienced as runoff and landscape drainage transport surface water to landscapes distal to the precipitation events[67]. Moreover, a more distal influence of precipitation would also fit with the strong associations between outbreaks and both riparian systems and hydrological flow accumulation. Conversely, it may be that the primary influence of La Niña-associated anomalies on JEV risk operates more indirectly with respect to the distribution of wild hosts across the landscape rather than reflecting a direct impact on mosquito ecology. For example, evidence suggests that La Niña phase increases in precipitation increase breeding among many species of birds, including inland waterbirds, in temperate Australia[68]. Specifically, La Nina anomalies were associated with both an earlier start to the breeding season and a longer breeding period in temperate, but not arid, Australia, which would both increase the number and period of availability of JEV susceptible bird hosts.

These important questions regarding altered weather patterns cannot be answered with the current data for the following reasons. First, as mentioned, JEV risk cannot be assessed at sufficiently local spatial scale, because outbreaks were not reported at sufficiently local scale. Second, the temporal granularity is necessarily coarse since there exist only a limited number of outbreaks within only a single season of observation. Notwithstanding the fact that the observed La Niña-associated anomalies in 2021 preceded the JEV outbreaks reported early in 2022, the correct temporal direction does not imply causation. Moreover, as mentioned above, the temporal direction of the association, even if genuinely causal, does not necessarily indicate whether the anomalies operate directly with respect to mosquito ecology or indirectly with respect to waterbird host ecology. In summary, the current data are limited in both spatial and temporal scale thus impeding the ability to infer causal relationships between altered weather patterns and JEV outbreaks.

Beyond the limitations already discussed, additional comment on this study’s further limitations is warranted. First, given that the unique geographic emergence of JEV in 2022 is the first of its kind in Australia, the current work is based on a small sample size and as such the risk estimates were associated with considerable uncertainty as quantified in the confidence limits provided. As an additional assessment of the impact of this uncertainty on model utility, a sensitivity analysis was conducted assessing model performance at the upper and lower confidence limits. This showed that despite a reduction in performance at the lower confidence limit, model outputs still performed reasonably well and thus still demonstrated utility across the spectrum of uncertainty. Second, reporting bias may be present despite diligent reporting of outbreaks to WOAH. As such, background points were sampled proportional to mean pig density as an indicator of disease detection likelihood to correct for potential reporting bias. Third, the ardeid species distribution models relied on human observations of birds and therefore are also subject to bias, because bird accessibility may influence reporting effort. Therefore, reporting bias in bird observations was corrected by sampling background points proportional to HFP. Fourth, while this study was able to estimate the landscape suitability of all extant Australian ardeid species, we also note that suitability is a representation of the fundamental niche only as reflected by the abiotic environmental features used to model their distributions. Conversely, the realised niche of any given species must also be determined by biotic interactions at the level of the local community and by the dispersal ability and history of a particular species relative to a local area given that environmental filtering is favourable to the species. Critically, neither the spatial nor temporal granularity of the current study allowed evaluation of interspecific interaction or dispersal history. Moreover, biotic interactions, and their subsequent effects on community composition, may have been considerably altered following the exceptional climate anomalies associated with the 2021 La Niña. It is therefore worth reiterating that the metric of ardeid species richness described here is not intended as a description of the actual community composition at local scale, but rather as the potential community assemblage given favourable environmental filtering alone. Importantly, an accounting of interspecific interaction should be incorporated into future work, including surveillance mechanisms, that seeks to more comprehensively evaluate the infection ecology of JEV in Australia.

In conclusion, this investigation has provided a preliminary account of the landscape epidemiology of emergent JEV in eastern Australia, thereby demarcating risk across heterogeneous landscapes. Accordingly, risk was delineated primarily by ardeid habitat within mosaics of riparian wetlands and agricultural land use, and the increasing accumulation of surface water runoff in the landscape. While preliminary, these findings highlight the importance of water presence in, and movement through, landscape patches comprising ardeid habitat and cultivated land. This suggests the importance of incorporating wild waterbird surveillance into ongoing monitoring systems that are currently surveying mosquitoes and piggeries in affected areas, as well as expanding surveillance to include locations that are embedded within key anthropogenic ecotones and exhibit a propensity to flooding. It is also worth noting that, while the results of this study primarily describe the landscape of piggery outbreak risk, some environmental features may also share important structural elements with human outbreak risk, such as ardeid habitat suitability and the movement of water through and accumulation in the landscape. Nevertheless, we must also consider that there may be multiple landscapes favourable to the circulation of JEV across eastern Australia. Although the 2022 JEV outbreaks in piggeries did not demonstrate an association with the distribution of feral pigs, for example, distinct foci of infection among feral pigs may nevertheless currently exist, or may emerge in the future, in landscapes quite distinct from those delineating the widespread emergence in domestic pigs.

## Acknowledgements

We thank SunPork Group for information about case farms, and Richard Bradhurst from the University of Melbourne who facilitated the sharing of the data products.

## Data Availability Statement

All data described in this study are publicly available for download at the cited sources in the text except the pig density data, which contains sensitive information to the Australian agricultural industry (e.g. locations of piggeries). Provision of this data for third parties requires their application to Australian Pork Limited.

## References

[1] J.S. Mackenzie, D.T. Williams, A.F. van den Hurk, D.W. Smith, B.J. Currie, Japanese Encephalitis Virus: The Emergence of Genotype IV in Australia and Its Potential Endemicity, Viruses. 14 (2022) 2480. https://doi.org/10.3390/V14112480.

[2] Australian Department of Health and Aged Care, Japanese encephalitis virus situation declared a Communicable Disease Incident of National Significance | Australian Government Department of Health and Aged Care, (2022). https://www.health.gov.au/news/japanese-encephalitis-virus-situation-declared-a-communicable-disease-incident-of-national-significance (accessed November 4, 2022).

[3] C.R. Williams, C.E. Webb, S. Higgs, A.F. van den Hurk, Japanese Encephalitis Virus Emergence in Australia: Public Health Importance and Implications for Future Surveillance, Vector Borne Zoonotic Dis. 22 (2022) 529–534. https://doi.org/10.1089/VBZ.2022.0037.

[4] J.N. Hanna, S.A. Ritchie, D.A. Phillips, J. Shield, M.C. Bailey, J.S. Mackenzie, M. Poidinger, B.J. McCall, P.J. Mills, An outbreak of Japanese encephalitis in the Torres Strait, Australia, 1995, Med J Aust. 165 (1996) 256–260. https://doi.org/10.5694/J.1326-5377.1996.TB124960.X.

[5] J.N. Hanna, S.A. Ritchie, D.A. Phillips, J.M. Lee, S.L. Hills, A.F. van den Hurk, A.T. Pyke, C.A. Johansen, J.S. Mackenzie, Japanese encephalitis in north Queensland, Australia, 1998, Med J Aust. 170 (1999) 533–536. https://doi.org/10.5694/J.1326-5377.1999.TB127878.X.

[6] A.F. van den Hurk, A.T. Pyke, J.S. Mackenzie, S. Hall-Mendelin, S.A. Ritchie, Japanese Encephalitis Virus in Australia: From Known Known to Known Unknown, Trop Med Infect Dis. 4 (2019). https://doi.org/10.3390/TROPICALMED4010038.

[7] World Health Organization, Japanese encephalitis, Fact Sheets: Japanese Encephalitis. (2019). https://www.who.int/news-room/fact-sheets/detail/japanese-encephalitis (accessed November 25, 2022).

[8] Y. Cheng, N.T. Minh, Q.T. Minh, S. Khandelwal, H.E. Clapham, Estimates of Japanese Encephalitis mortality and morbidity: A systematic review and modeling analysis, PLoS Negl Trop Dis. 16 (2022) e0010361. https://doi.org/10.1371/JOURNAL.PNTD.0010361.

[9] World Health Organization, Japanese encephalitis vaccines: WHO position paper, Weekly Epidemiological Record. 90 (2015) 69–87.

[10] F.M. Rodrigues, S.N. Guttikar, B.D. Pinto, Prevalence of antibodies to Japanese encephalitis and West Nile viruses among wild birds in the Krishna-Godavari Delta, Andhra Pradesh, India, Trans R Soc Trop Med Hyg. 75 (1981) 258–262. https://doi.org/10.1016/0035-9203(81)90330-8.

[11] A. v. Jamgaonkar, P.N. Yergolkar, G. Geevarghese, G.D. Joshi, M. v. Joshi, A.C. Mishra, Serological evidence for Japanese encephalitis virus and West Nile virus infections in water frequenting and terrestrial wild birds in Kolar District, Karnataka State, India. A retrospective study, Acta Virol. 47 (2003) 185–188.

[12] E.L. Buescher, W.F. Scherer, M.Z. Rosenberg, L.J. Kutner, Immunologic studies of Japanese encephalitis virus in Japan. IV. Maternal antibody in birds., Journal of Immunology. 83 (1959) 614–619.

[13] S. Bhattacharya, P. Basu, Japanese Encephalitis Virus (JEV) infection in different vertebrates and its epidemiological significance: a Review, International Journal of Fauna and Biological Studies. 1 (2014) 32–37. https://pdfs.semanticscholar.org/51c4/860d8c66c5fa9d730143e836b8db0b21c1a4.pdf.

[14] A. Baruah, R. Hazarika, N. Barman, S. Islam, B. Gulati, Mosquito abundance and pig seropositivity as a correlate of Japanese encephalitis in human population in Assam, India, J Vector Borne Dis. 55 (2018) 291–296. https://doi.org/10.4103/0972-9062.256564.

[15] J. Borah, P. Dutta, S.A. Khan, J. Mahanta, Epidemiological concordance of Japanese encephalitis virus infection among mosquito vectors, amplifying hosts and humans in India, Epidemiol Infect. 141 (2013) 74–80. https://doi.org/10.1017/S0950268812000258.

[16] M. Kakkar, S. Chaturvedi, V.K. Saxena, T.N. Dhole, A. Kumar, E.T. Rogawski, S. Abbas, V. v. Venkataramanan, P. Chatterjee, Identifying sources, pathways and risk drivers in ecosystems of Japanese Encephalitis in an epidemic-prone north Indian district, PLoS One. 12 (2017) 1– 17. https://doi.org/10.1371/journal.pone.0175745.

[17] B. Chen, B. Beaty, Japanese encephalitis vaccine (2-8 strain) and parent (SA 14 strain) viruses in Culex tritaeniorhynchus mosquitoes, American Journal of Tropical Medicine and Hygiene. 31 (1982) 403–407.

[18] K. Komada, N. Sasaki, Y. Inoue, Studies of live attenuated Japanese encephalitis vaccine in swine, Journal of Immunology. 100 (1968) 194–200.

[19] S. Ghimire, S. Dhakal, N.P. Ghimire, D.D. Joshi, Pig Sero-Survey and Farm Level Risk Factor Assessment for Japanese Encephalitis in Nepal, Int J Appl Sci Biotechnol. 2 (2014) 311–314. https://doi.org/10.3126/ijasbt.v2i3.10639.

[20] A.F. van den Hurk, D.J. Nisbet, R.A. Hall, B.H. Kay, J.S. Mackenzie, S.A. Ritchie, Vector competence of Australian mosquitoes (Diptera: Culicidae) for Japanese encephalitis virus, J Med Entomol. 40 (2003) 82–90. https://doi.org/10.1603/0022-2585-40.1.82.

[21] A.F. van den Hurk, E. Skinner, S.A. Ritchie, J.S. Mackenzie, The Emergence of Japanese Encephalitis Virus in Australia in 2022: Existing Knowledge of Mosquito Vectors, Viruses. 14 (2022). https://doi.org/10.3390/V14061208.

[22] M. Furlong, A. Adamu, R.I. Hickson, P. Horwood, M. Golchin, A. Hoskins, T. Russell, Estimating the Distribution of Japanese Encephalitis Vectors in Australia Using Ecological Niche Modelling, Tropical Medicine and Infectious Disease 2022, Vol. 7, Page 393. 7 (2022) 393. https://doi.org/10.3390/TROPICALMED7120393.

[23] M.G. Walsh, A. Pattanaik, N. Vyas, D. Saxena, C. Webb, S. Sawleshwarkar, C. Mukhopadhyay, High-risk landscapes of Japanese encephalitis virus outbreaks in India converge on wetlands, rain-fed agriculture, wild Ardeidae, and domestic pigs and chickens, Int J Epidemiol. (2022). https://doi.org/10.1093/ije/dyac050.

[24] M.J. McPhaden, El Niño and La Niña: Causes and global consequences, Encyclopedia of Global Environmental Change. 1 (2001) 353–370.

[25] A. Bureau of Meteorology, Recent and historical rainfall maps, Australian Bureau of Meteorology, Recent and Historical Rainfall Maps. (2021). http://www.bom.gov.au/climate/maps/rainfall/?variable=rainfall&map=totals&period=12month&region=nat&year=2021&month=12&day=31 (accessed November 9, 2022).

[26] N.A. Prow, The changing epidemiology of Kunjin virus in Australia, Int J Environ Res Public Health. 10 (2013) 6255–6272. https://doi.org/10.3390/IJERPH10126255.

[27] L.A. Selvey, L. Dailey, M. Lindsay, P. Armstrong, S. Tobin, A.P. Koehler, P.G. Markey, D.W. Smith, The changing epidemiology of Murray Valley encephalitis in Australia: the 2011 outbreak and a review of the literature, PLoS Negl Trop Dis. 8 (2014) 18. https://doi.org/10.1371/JOURNAL.PNTD.0002656.

[28] G. of I. National vectorborne Disease Control Program, Ministry of Health and Family Welfare, Statewise number of AES/JE cases and deaths from 2010–2017., (2017).

[29] World Organisation For Animal Health, World Animal Health Information System (WAHIS), WAHIS Inicident Reports. (2022). https://wahis.woah.org/#/home (accessed November 4, 2022).

[30] New South Wales Health, Surveillance and monitoring weekly reports season 2021-22 - Vector-borne diseases, Vector-Borne Diseases Surveillance and Monitoring Weekly Reports. (2022). https://www.health.nsw.gov.au/environment/pests/vector/Pages/nswasp-weekly-report-2021-22.aspx (accessed December 18, 2022).

[31] N. McKilligan, Herons, Egrets and Bitterns: Their Biology and Conservation in Australia, CSIRO Publishing, 2005. https://www.publish.csiro.au/book/4841 (accessed November 4, 2022).

[32] Global Biodiversity Information Facility, GBIF occurrence download—Ardeidae Australia, Global Biodiversity Information Facility. (2022). https://doi.org/10.15468/dl.6fsjpk (accessed November 4, 2022).

[33] Global Biodiversity Information Facility, GBIF Occurrence Download - Sus scrofa, Global Biodiversity Information Facility. (2022). https://doi.org/10.15468/dl.apcrgq (accessed November 4, 2022).

[34] R.A. Bradhurst, S.E. Roche, I.J. East, P. Kwan, M. Graeme Garner, A hybrid modeling approach to simulating foot-and-mouth disease outbreaks in Australian livestock, Front Environ Sci. 3 (2015) 17. https://doi.org/10.3389/FENVS.2015.00017/BIBTEX.

[35] E.W. Sanderson, M. Jaiteh, M.A. Levy, K.H. Redford, A. v. Wannebo, G. Woolmer, The Human Footprint and the Last of the Wild, Bioscience. 52 (2002) 891. https://doi.org/10.1641/0006-3568(2002)052[0891:THFATL]2.0.CO;2.

[36] Socioeconomic Data and Applications Center | SEDAC, Methodslll" Last of the Wild, v2 | SEDAC, (n.d.). http://sedac.ciesin.columbia.edu/data/collection/wildareas-v2/methods (accessed December 23, 2014).

[37] C. Lamarche, M. Santoro, S. Bontemps, R. d’Andrimont, J. Radoux, L. Giustarini, C. Brockmann, J. Wevers, P. Defourny, O. Arino, Compilation and Validation of SAR and Optical Data Products for a Complete and Global Map of Inland/Ocean Water Tailored to the Climate Modeling Community, Remote Sensing 2017, Vol. 9, Page 36. 9 (2017) 36. https://doi.org/10.3390/RS9010036.

[38] European Space Agency, Climate Change Initiative, Land Cover project, 2017: Water Bodies v4.0, ESA (European Space Agency) CCI (Climate Change Initiative) Land Cover Project 2017. (2017). http://maps.elie.ucl.ac.be/CCI/viewer/ (accessed November 4, 2022).

[39] European Space Agency, Climate Change Initiative, Land Cover CCI Product - Annual LC maps from 2000 to 2015 (v2.0.7)., ESA (European Space Agency) CCI (Climate Change Initiative) Land Cover Project 2017. (2017). http://maps.elie.ucl.ac.be/CCI/viewer/ (accessed November 4, 2022).

[40] S. of G. and E.S. University of Southampton, D. of G. and G. University of Louisville, D. de G. Universite de Namur, WorldPop hub, (n.d.). https://hub.worldpop.org/doi/10.5258/SOTON/WP00644 (accessed November 4, 2022).

[41] WorldPop, WorldPoplll:: Geospatial covariate data layers, (n.d.). https://hub.worldpop.org/project/categories?id=14 (accessed December 2, 2022).

[42] B. Lehner, K. Verdin, A. Jarvis, HydroSHEDS Technical Documentation, 2006. http://hydrosheds.cr.usgs.gov.

[43] A. McNally, FLDAS Noah Land Surface Model L4 Global Monthly Anomaly 0.1 × 0.1 degree (MERRA-2 and CHIRPS), Goddard Earth Sciences Data and Information Services Center (GES DISC). (2018). https://disc.gsfc.nasa.gov/datasets/FLDAS_NOAH01_C_GL_MA_001/summary (accessed November 4, 2022).

[44] S.E. Fick, R.J. Hijmans, WorldClim 2: new 1-km spatial resolution climate surfaces for global land areas, International Journal of Climatology. (2017). https://doi.org/10.1002/joc.5086.

[45] C.H.B. Priestley, R.J. Taylor, On the assessment of surface heat flux and evaporation using large-scale parameters, Mon Weather Rev. 100 (1972) 81–92. https://doi.org/10.1175/1520-0493(1972)100<0081:OTAOSH>2.3.CO;2.

[46] a. Khaldi, A. Khaldi, A. Hamimed, Using the Priestley-Taylor expression for estimating actual evapotranspiration from satellite Landsat ETM + data, Proceedings of the International Association of Hydrological Sciences. 364 (2014) 398–403. https://doi.org/10.5194/piahs-364-398-2014.

[47] A. Trabucco, R.J. Zomer, Global soil water balance geospatial database, CGIAR Consortium for Spatial Information. (2010). http://www.cgiar-csi.org.

[48] J. Elith, J.R. Leathwick, T. Hastie, A working guide to boosted regression trees., J Anim Ecol. 77 (2008) 802–13. https://doi.org/10.1111/j.1365-2656.2008.01390.x.

[49] J. Friedman, Greedy function approximation: a gradient boosting machine, Ann Stat. 29 (2001) 1189–1232.

[50] L. Breiman, Random forests, Mach Learn. 45 (2001) 5–32. https://doi.org/10.1023/A:1010933404324.

[51] G. James, D. Witten, T. Hastie, R. Tibshirani, An introduction to Statistical Learning, 2000. https://doi.org/10.1007/978-1-4614-7138-7.

[52] S.N. Wood, Stable and Efficient Multiple Smoothing Parameter Estimation for Generalized Additive Models, J Am Stat Assoc. 99 (2004) 673–686. https://doi.org/10.1198/016214504000000980.

[53] S.N. Wood, Generalized additive modelslll: an introduction with R, 2nd ed., Chapman and Hall/CRC, New York, 2017. https://books.google.com.au/books?id=HL-PDwAAQBAJ&printsec=frontcover&dq=Wood,+S.N.+(2017)+Generalized+Additive+Models:+An+Introduction+with+R+(2nd+edition).+Chapman+and+Hall/CRC.&hl=en&sa=X&ved=0ahUKEwiB_43x9NvjAhUC4nMBHU7CDWQQ6AEIMDAB#v=onepage&q=Wood%2 (accessed July 30, 2019).

[54] B. Naimi, M.B. Araújo, sdm: a reproducible and extensible R platform for species distribution modelling, Ecography. 39 (2016) 368–375. https://doi.org/10.1111/ecog.01881.

[55] G.G. Mittelbach, B.J. McGill, Community Ecology, Second, Oxford University Press, 2019.

[56] R Core Team, R: A language and environment for statistical computing v. 4.1.2, Vienna, 2022. http://www.R-project.org/.

[57] A. Baddeley, R. Turner, Practical Maximum Pseudolikelihood for Spatial Point Patterns (with Discussion), Aust N Z J Stat. 42 (2000) 283–322. https://doi.org/10.1111/1467-842X.00128.

[58] A. Baddeley, R. Turner, spatstat: An R Package for Analyzing Spatial Point Patterns, Journal of Statistical Software 12(6). (2005). http://www.jstatsoft.org/v12/i06/ (accessed October 23, 2014).

[59] A. Baddeley, E. Rubak, R. Turner, Spatial Point Patterns: Methodology and Applications with R, CRC Press, 2015. https://books.google.com/books?id=rGbmCgAAQBAJ&pgis=1 (accessed February 5, 2016).

[60] D.J. Civitello, J. Cohen, H. Fatima, N.T. Halstead, J. Liriano, T.A. McMahon, C.N. Ortega, E.L. Sauer, T. Sehgal, S. Young, J.R. Rohr, Biodiversity inhibits parasites: Broad evidence for the dilution effect, Proceedings of the National Academy of Sciences. 112 (2015) 8667–8671. https://doi.org/10.1073/pnas.1506279112.

[61] W. Scherer, Ecological studies of Japanese encephalitis in Japan. Parts I-IX, American Journal of Tropical Medicine and Hygiene. 8 (1959) 644–722.

[62] M. Ogata, Y. Nagao, F. Jitsunari, N. Kitamura, T. Okazaki, Infection of herons and domestic fowls with Japanese encephalitis virus with specific reference to maternal antibody of hen (epidemiological study on Japanese encephalitis 26), Acta Med Okayama. 24 (1970) 175–184.

[63] N. Nemeth, A. Bosco-Lauth, P. Oesterle, D. Kohler, R. Bowen, North American birds as potential amplifying hosts of Japanese encephalitis virus, American Journal of Tropical Medicine and Hygiene. 87 (2012) 760–767. https://doi.org/10.4269/ajtmh.2012.12-0141.

[64] C.E. Webb, S.L. Doggett, R.C. Russell, A guide to the mosquitoes of Australia, First, CSIRO Publishing, 2016.

[65] J.K. Hanford, C.E. Webb, D.F. Hochuli, Habitat Traits Associated with Mosquito Risk and Aquatic Diversity in Urban Wetlands, Wetlands 2019 39:4. 39 (2019) 743–758. https://doi.org/10.1007/S13157-019-01133-2.

[66] J.K. Hanford, C.E. Webb, D.F. Hochuli, Management of urban wetlands for conservation can reduce aquatic biodiversity and increase mosquito risk, Journal of Applied Ecology. 57 (2020) 794–805. https://doi.org/10.1111/1365-2664.13576.

[67] D.C. Verdon, A.M. Wyatt, A.S. Kiem, S.W. Franks, Multidecadal variability of rainfall and streamflow: Eastern Australia, Water Resour Res. 40 (2004) 10201. https://doi.org/10.1029/2004WR003234.

[68] D. Englert Duursma, R. v. Gallagher, S.C. Griffith, Effects of El Niño Southern Oscillation on avian breeding phenology, Divers Distrib. 24 (2018) 1061–1071. https://doi.org/10.1111/DDI.12750.

