## Supplemental material for "Little pig, little pig, let me come in: The influence of landscape structure and La Niña climatic anomalies on the emergence of Japanese encephalitis virus in Australian piggeries in 2022"

Table S1. Ardeidae species comparisons based on ensemble species distribution models. Each species listed is presented with their associated number of observations in the field (and the number of observations used for analysis after thinning in parentheses), their suitability model fit (deviance), and model performance (area under the receiver operating characteristic curve [AUC]).

| **Ardeidae species** | **Number of field observations** | **Deviance** | **AUC (%)** |
|---|---|---|---|
| *Ardea alba* | 14957 (9771) | 0.70 | 92 |
| *Ardea modesta** | 9842 (5682) | 0.67 | 93 |
| *Ardea pacifica* | 19201 (14666) | 0.95 | 86 |
| *Ardea sumatrana* | 293 (165) | 0.23 | 100 |
| *Botaurus poiciloptilus* | 644 (466) | 0.65 | 94 |
| *Bubulcus coromandus* | 142 (131) | 0.41 | 98 |
| *Bubulcus ibis** | 14419 (9477) | 0.58 | 95 |
| *Butorides striata* | 3796 (2192) | 0.28 | 99 |
| *Dupetor flavicollis* (*Ixobrychus flavicollis*) | 715 (570) | 0.52 | 96 |
| *Egretta garzetta* | 9249 (5204) | 0.56 | 95 |
| *Egretta intermedia* (*Ardea intermedia*) | 9449 (5809) | 0.68 | 93 |
| *Egretta novaehollandiae* | 48375 (27728) | 0.83 | 89 |
| *Egretta picata* | 368 (260) | 0.56 | 95 |
| *Egretta sacra* | 2130 (1338) | 0.24 | 99 |
| *Ixobrychus dubius* | 378 (255) | 0.52 | 97 |
| *Nycticorax caledonicus* | 6987 (4655) | 0.84 | 89 |

* *Ardea modesta* was previously considered a subspecies of *Ardea alba* and *Bubulcus coromandus* was previously considered a subspecies of *Bubulcus ibis*. These were ultimately combined to estimate *Ardea alba* and *Bubulcus coromandus* habitat suitabilities, respectively.

Table S2. Crude, univariable regression coefficients and 95% confidence intervals for the associations between Japanese encephalitis virus outbreaks and each landscape feature as derived from an inhomogeneous Poisson model.

| **Landscape feature** | **AIC** | **Coefficient** | **95% confidence interval** |
|---|---|---|---|
| Null model | 206.95 | | |
| **Climate** | | | |
| Mean annual precipitation above the climate average (mm/day) | 206.50 | -0.005 | -0.11 – 0.097 |
| Mean June & Sept precipitation above the climate average (mm/day) | 170.09 | 0.32 | 0.20 – 0.45 |
| Mean annual temperature below the climate average (Celsius) | 196.24 | 0.20 | 0.08 – 0.32 |
| Mean annual soil moisture above the climate average (10 cm depth) | 204.77 | -13.50 | -36.72 – 9.72 |
| **Hydrogeography and surface hydrology** | | | |
| Distance to inland wetlands (1 km) | 193.49 | -0.04 | -0.07 – -0.02 |
| Distance to rivers and streams (1 km) | 157.83 | -0.14 | -0.21 – -0.07 |
| Hydrological flow accumulation (per 100 1 $km^2$ land parcels of accumulation) | 195.76 | 0.003 | 0.002 – 0.004 |
| **Land cover and land use** | | | |
| Distance to forest (1 km) | 196.46 | -0.04 | -0.07 – -0.005 |
| Distance to shrubland (1 km) | 191.60 | -0.10 | -0.19 – -0.02 |
| Distance to grassland (1 km) | 197.28 | -0.05 | -0.09 – -0.01 |
| Distance to aquatic vegetation (1 km) | 208.17 | -0.005 | -0.02 – 0.008 |
| Distance to cultivated land (1 km) | 132.09 | -0.22 | -0.29 – -0.15 |
| **Animal hosts** | | | |
| Ardeidae richness (N) | 171.17 | 1.96 | 1.18 – 2.73 |
| Feral pig habitat suitability (%) | 196.17 | -2.90 | -4.69 – -1.10 |

Figure S1. Mean monthly above average precipitation anomaly during 2021.

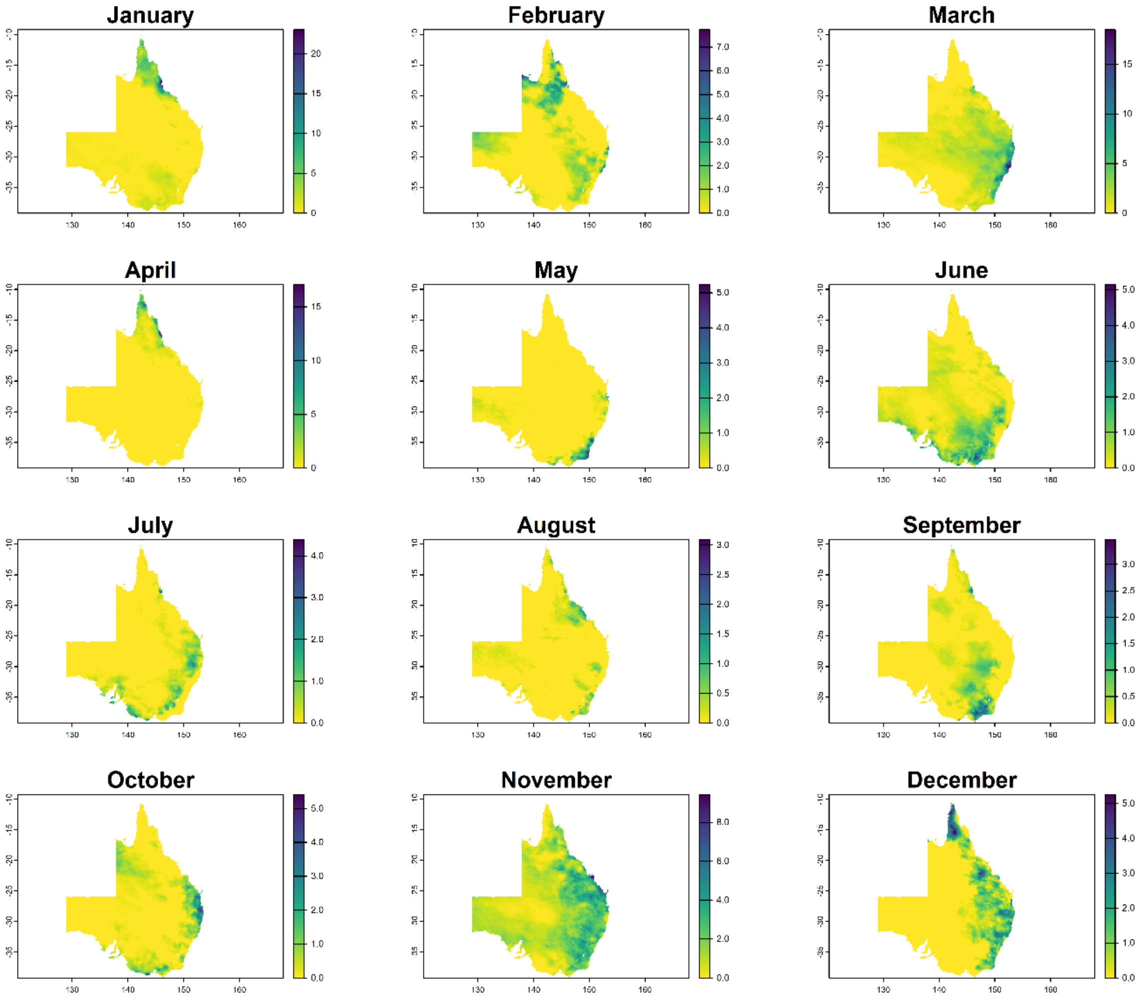

Table S3. Crude, univariable regression coefficients and 95% confidence intervals for the associations between Japanese encephalitis virus outbreaks and the mean above average precipitation anomaly for each month during 2021 as derived from an inhomogeneous Poisson model.

| Mean above average precipitation anomaly | AIC | Coefficient | 95% confidence interval |
|---|---|---|---|
| January | 206.24 | -0.05 | -0.27 – 0.16 |
| February | 200.49 | -0.62 | -1.22 – -0.02 |
| March | 205.03 | -0.08 | -0.22 – 0.06 |
| April | 194.67 | -7.33 | -17.89 – 3.22 |
| May | 191.73 | -7.69 | -16.02 – 0.65 |
| June | 185.59 | 0.76 | 0.48 – 1.04 |
| July | 204.73 | 0.44 | -0.16 – 1.05 |
| August | 194.81 | -7.81 | -15.01 – -0.60 |
| September | 202.45 | 0.62 | 0.08 – 1.15 |
| October | 205.86 | -0.17 | -0.62 – 0.27 |
| November | 206.27 | 0.05 | -0.15 – 0.25 |
| December | 200.80 | -0.51 | -1.00 – -0.03 |

Figure S2. La Niña precipitation and temperature anomalies during 2021 and hydrological flow accumulation. Salmon-coloured points indicate locations of Japanese encephalitis outbreaks in piggeries (data publicly available from the World Animal Health Information System; https://www.woah.org/en/what-we-do/animal-health-and-welfare/disease-data-collection/world-animal-health-information-system/, accessed 19 December 2022).

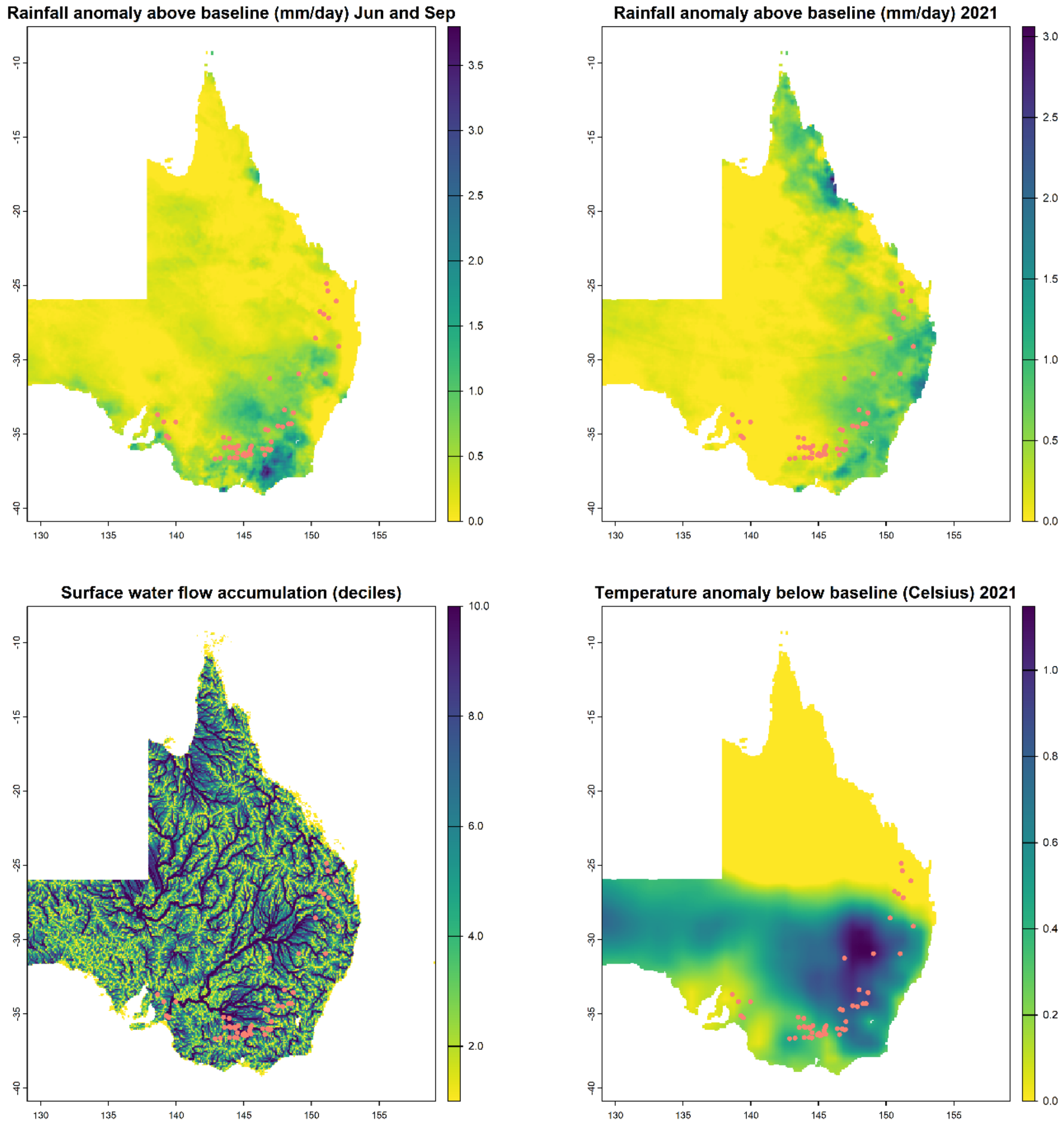

Figure S3. Proximity to wetland and land cover patches and Ardeidae richness. Blue-coloured points indicate locations of Japanese encephalitis outbreaks in piggeries (data publicly available from the World Animal Health Information System; https://www.woah.org/en/what-we-do/animal-health-and-welfare/disease-data-collection/world-animal-health-information-system/, accessed 19 December 2022).

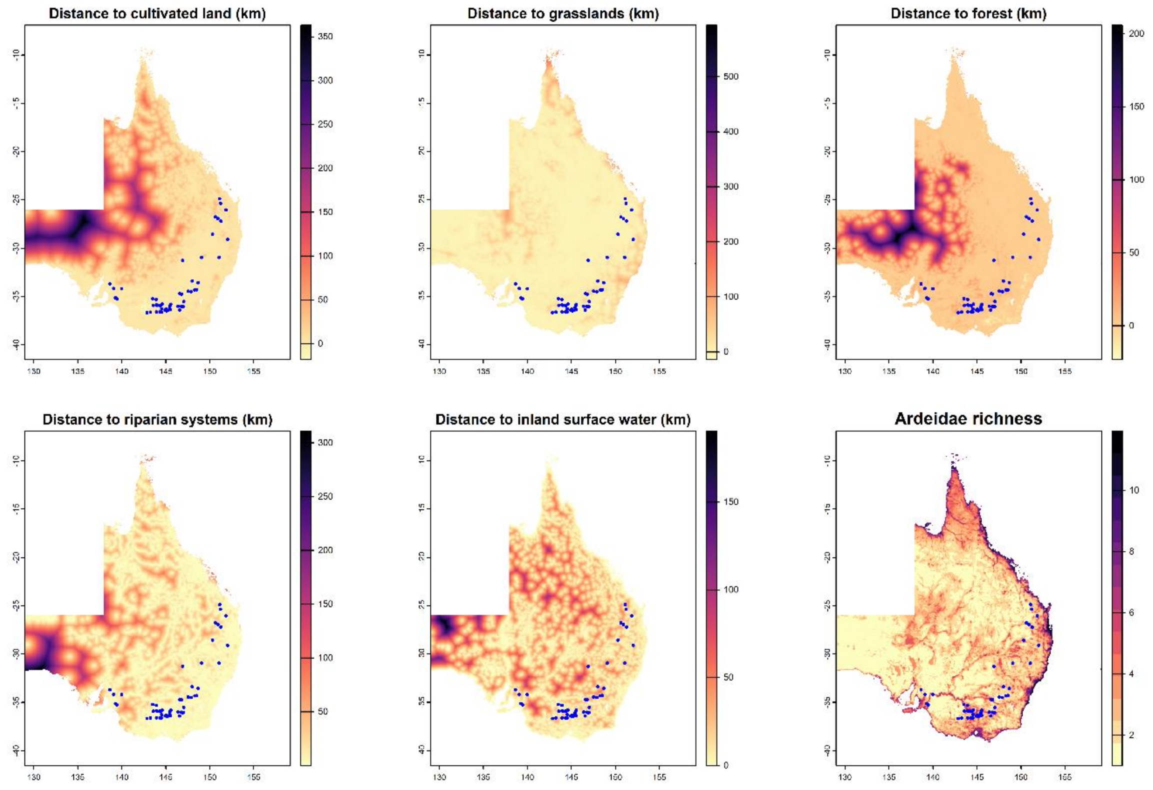

Table S4. Japanese encephalitis virus (JEV) outbreak multiple inhomogeneous Poisson process model comparisons by Akaike information criterion (AIC) and area under the receiver operating characteristic curve (AUC). Each nested multiple point process model includes those variables that were univariably associated with JEV outbreaks (Table S2).

| Pont process models | AIC | AUC (%) |
|---|---|---|
| **Model 1 (Climate only)**: *Mean above average precipitation anomaly + Mean below average temperature anomaly* | 187.47 | 85.4 |
| **Model 2 (Wetlands and hydrology):** *Inland wetland proximity + river proximity + hydrological flow accumulation* | 151.45 | 88.8 |
| **Model 3 (land cover/land use)**: *Cultivated land proximity + grassland proximity + forest proximity* | 121.81 | 95.2 |
| **Model 4 (Reservoir hosts only)**: *Ardeidae richness + Ardeidae richness $^2$* | 171.17 | 90.4 |
| **Model 5 (Full)**: *Ardeidae richness + Ardeidae richness $^2$ + Inland wetland proximity + river proximity + hydrological flow accumulation + cultivated land proximity + grassland proximity + forest proximity + mean above average precipitation anomaly + mean below average temperature anomaly* | 92.20 | 93.3 |
| **Model 6 (Final)**: *Ardeidae richness + Ardeidae richness $^2$ + river proximity + cultivated land proximity + grassland proximity + hydrological flow accumulation* | 91.65 | 93.6 |
| **Model 7 (Final, with river:cultivation interaction)**: *Ardeidae richness + Ardeidae richness $^2$ + river proximity + cultivated land proximity + river:cultivation interaction + grassland proximity + hydrological flow accumulation* | 93.01 | 93.6 |
